# Cap-independent translation and a precisely localized RNA sequence enable SARS-CoV-2 to control host translation and escape anti-viral response

**DOI:** 10.1101/2021.08.18.456855

**Authors:** Boris Slobodin, Urmila Sehrawat, Anastasia Lev, Ariel Ogran, Davide Fraticelli, Daniel Hayat, Binyamin Zuckerman, Igor Ulitsky, Amir Ben-Shmuel, Elad Bar-David, Haim Levy, Rivka Dikstein

## Abstract

Translation of SARS-CoV-2-encoded mRNAs by the host ribosomes is essential for its propagation. Following infection, the early expressed viral protein NSP1 binds the ribosome, represseses translation and induces mRNA degradation, while the host elicits anti-viral response. The mechanisms enabling viral mRNAs to escape this multifaceted repression remain obscure. Here we show that expression of NSP1 leads to destabilization of multi-exon cellular mRNAs, while intron-less transcripts, such as viral mRNAs and anti-viral interferon genes, remain relatively stable. We identified a conserved and precisely located cap-proximal RNA element devoid of guanosines that confers resistance to NSP1-meidated translation inhibition. Importantly, the primary sequence rather than the secondary structure is critical for protection. We further show that the genomic 5’UTR of SARS-CoV-2 exhibits an IRES-like activity and promotes expression of NSP1 in an eIF4E-independent and Torin-1 resistant manner. Upon expression, NSP1 enhances cap-independent translation. However, the sub-genomic 5’UTRs are highly sensitive to eIF4E availability, rendering viral propagation partially sensitive to Torin-1. The combined NSP1-mediated degradation of spliced mRNAs and translation inhibition of single-exon genes, along with the unique features present in the viral 5’UTRs, ensure robust expression of viral mRNAs. These features can be exploited as potential therapeutic targets.

## Introduction

Viruses do not encode functional translation machinery and rely on the host cell to translate their genetic information into proteins. This makes viruses obligatory parasites that must exploit the host translation apparatus for successful proliferation. Such an absolute dependence turns the translational regulation into the Achilles heel of viral infection and propagation. Infected cells attenuate cap-dependent translation as means of preventing translation of viral proteins (1). Translation that does not rely on the 5’cap structure, such as via use of internal ribosome entry sites (IRESes), is a major strategy employed by multiple positive single-stranded RNA viruses to support expression of viral mRNAs when the cap-dependent translation is inhibited. Although IRES-mediated translation initiation is less efficient than the canonical cap-dependent mode, it allows eIF4E-independent translation and, therefore, is particularly important upon mTOR inhibition (2). In parallel, multiple viruses attempt to enforce mTOR activation to relief the translational repression (3). Notably, the cap-dependent and -independent pathways of translation initiation are not mutually exclusive and may cooperate to enhance viral-specific translational yield, as was shown for HCV (4).

SARS-CoV-2 is a betacoronavirus encoding a long 5’-capped and polyadenylated positive single-stranded RNA genome that serves as an immediate mRNA template for synthesis of ORF1ab, two precursor polyproteins that are proteolytically cleaved to form 16 non-structural proteins (5). In addition, the SARS-CoV-2 genome gives rise to multiple sub-genomic mRNAs that are synthesized via discontinuous RNA-templated transcription and, similarly to the genomic RNA, bear 5’caps and poly(A) tails (6). The presence of the 5’cap on all viral mRNAs stresses the importance of cap-dependent translation initiation for SARS-CoV-2 and raises the question regarding the ability of the virus to oppose its inhibition.

NSP1, the non-structural protein 1, is one of the major virulence factor encoded by SARS-CoV-2. NSP1 directly interacts with host ribosomes via its C’-terminal moiety (7, 8), repressing translation and inducing mRNA degradation in infected cells (9–11). Particularly, NSP1 was shown to associate with 40S ribosomal subunits at the mRNA entry channel, hindering the access of mRNAs to the ribosome. Interestingly, despite such a global effect, translation of innate immune genes is particularly repressed in the infected cells (10), enabling the virus to evade the host’s innate defense. Viral 5’UTRs were shown to protect mRNAs from NSP1-mediated inhibition (10, 12), and although the stem-loop element 1 (SL1) within viral 5’UTRs was suggested to play a role (13), the exact motif sufficient for protection remains insufficiently clear.

In this study, we show that NSP1 promotes degradation of host multi-exon mRNAs, while having a relatively minor effect on the stability of intron-less transcripts. Using single-exon reporter mRNAs, we demonstrate that these transcripts are repressed by NSP1 at the translation level. We identified a specific RNA sequence within the SARS-CoV-2 genomic and sub-genomic 5’UTRs, that enables viral mRNAs to escape repression. This element depends on its nucleotide composition rather than secondary structure and is characterized by the absence of guanosines and a precise location relative to the 5’end, two features that are conserved in several coronaviruses. We show that this element is sufficient for robust heterologous expression of DNA-encoded genes in NSP1-expressing cells. We also found that the genomic 5’UTR that drives NSP1 translation, exhibits an IRES-like activity, supporting eIF4E-independent, Torin-1 resistant translation initiation, while the sub-genomic 5’UTR is highly sensitive to eIF4E availability. This ability enables highly efficient expression of NSP1 even upon prolonged arrest of cap-dependent translation. These findings unravel the molecular strategies of SARS-CoV-2 to hijack and adjust the host translation machinery for viral propagation and to overcome both inhibition of cap-dependent translation and induction of interferons.

## Results

### NSP1 robustly destabilizes multi-exon mRNAs while moderately inhibiting translation initiation

It was recently reported that mRNAs of SARS-CoV-2-infected cells undergo massive destabilization (9, 10). In case of SARS-CoV-1, this role has been attributed to NSP1, the first protein encoded by the viral genome (18, 19). To test if NSP1 encoded by SARS-CoV-2 is sufficient to induce mRNA degradation, we transfected HEK293 cells with a plasmid encoding for NSP1 and isolated polyadenylated mRNAs after 24 hours of expression. Indeed, expression of NSP1 significantly reduced the levels of polyadenylated RNAs in cells (Fig. 1A). To directly examine the effect of NSP1 on mRNA stability, we measured half-lives of cellular mRNAs via time-course actinomycin D treatment followed by RNA-seq (20). Indeed, we found that expression of NSP1 strongly reduced the average half-life of mRNAs from 12 hours to 3.5 hours (Fig. 1B). Upon detailed mRNA analysis, we noticed that the magnitude of the effect on mRNA degradation differed according to the presence of introns: while the stability of multi-exon mRNAs was highly sensitive to NSP1, intron-less transcripts were relatively resistant (Fig. 1C). Using reporter genes with or without intron, we confirmed that the spliced gene is more sensitive to inhibition by NSP1 (Fig. 1D). Since viral mRNAs are intron-less, this observation provides a plausible explanation of how the virus ensures the relative stability of its own mRNAs. However, a major class of anti-viral genes, such as interferon alpha and beta, are all intron-less. It was reported that SARS-CoV-2 represses these genes mainly at the translation level (8), suggesting that the translation machinery of infected cells must distinguish, among intron-less transcripts, between viral and host mRNAs. Interestingly, long non-coding RNAs exhibited reduced vulnerability to NSP1-induced degradation (Fig. S1), indicating that NSP1-mediated RNA destabilization may coincide with recruitment to ribosomes. Overall, our results suggest that expression of NSP1 leads to global destabilization of multi-exon mRNAs, which may depend on the interaction with ribosomes.

**Figure 1:**
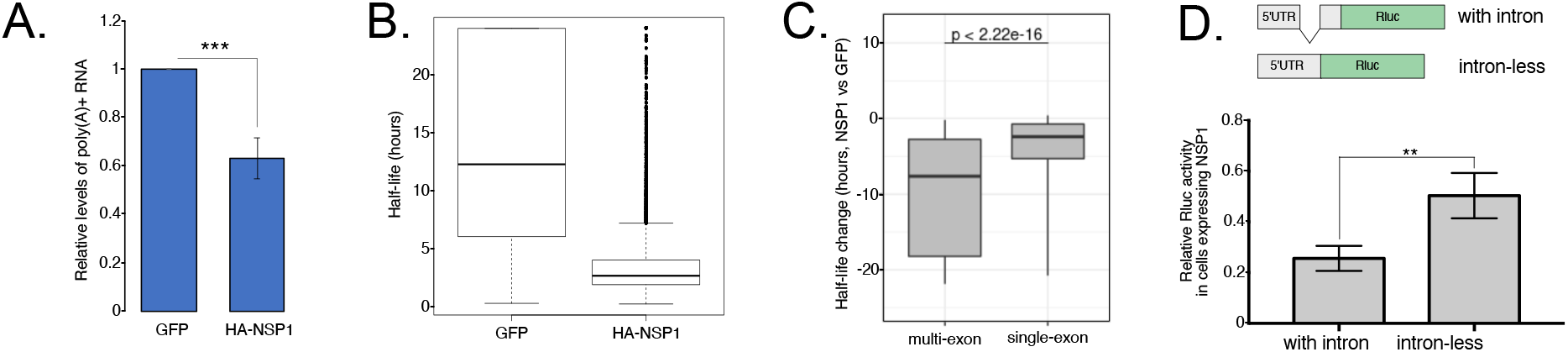
Expression of NSP1 destabilizes multi-exon mRNAs. **A.** HEK293 cells were transfected with plasmids encoding either eGFP or TSS-5’UTR-HA-NSP1 (4μg DNA per 10-cm dish) and collected after 24 hours. Isolated RNA was subjected to poly(A)-dependent fractionation, yields of which were quantified and plotted in a relative manner; n=5, bar represents SE. **B.** Cells transfected as detailed in (A) were treated with Actinomycin D (7.5μg/ml) for 0,2,4, and 6 hours, harvested and subjected to MARS-seq procedure to determine half-lives of polyadenylated RNAs; n=2. **C.** Changes in half-lives of multi- and single-exon mRNAs upon expression of HA-NSP1; n=2. **D.** HEK293 cells were transfected with plasmids encoding for Renilla luciferase (Rluc) reporter genes either with or without intron in the 5’UTR (upper panel) along with plasmids encoding for eGFP or HA-NSP1. After 24 hours, the relative expression of both reporters was calculated and plotted; n=3, bars represent SE.

NSP1 was reported to interact with the host ribosome near the entry channel and inhibit translation due to physical hindrance of the ribosome-mRNA interaction (7, 8, 21). To test the effect of NSP1 on cellular translation, we performed polysome profiling of HEK293 cells transfected with either HA-NSP1 or eGFP. Indeed, polysomal profiles of NSP1-expressing cells exhibited diminished polysomal fractions (Figs. 2A and S2C), indicating reduced translation. This conclusion was further supported by the puromycin incorporation assay. Puromycin is a structural analog of aminoacylated-tRNA (aa-tRNA), which leads to premature termination of translation, thus marking active translation. HEK293 cells were transfected with NSP1 or GFP for 48 hours and then subjected to a pulse of puromycin (10μg/ml) for 5 minutes and Western blotting using anti-puromycin antibodies. The results revealed a moderate loss of nascent polypeptide labeling upon NSP1 expression (Fig. 2B and S2A, B), confirming an attenuation of translation. Interestingly, we found that NSP1 protein was present only in the initiating fractions of the polysomal gradient and was devoid from the fractions of actively translated mRNAs (Fig. 2C). Taken together, these observations suggest that NSP1 may reduce the efficiency of the translation initiation step. As translation is linked to cell proliferation and survival, we examined cell growth after transfection of either NSP1 or GFP, anticipating that significantly reduced translation will negatively affect the proliferation ability of the cells. Remarkably, we did not detect any growth defects in the NSP1-expressing cells (Figs. 2D and S2D,E). To confirm these findings, we monitored cellular viability for up to 49 hours post-transfection with HA-NSP1 and compared it to the effect of known translational inhibitors, such as Torin1, puromycin, cycloheximide or homoharringtonine. We found that expression of NSP1 even slightly enhanced the growth kinetics, while application of low amounts of known translation inhibitors markedly reduced it (Figs. 2E and S2F). These observations uncover the relative extent of the NSP1 impact on cellular translation and mRNA stability and suggest that, at least within the tested time window, it does not interfere with cellular proliferation.

**Figure 2:**
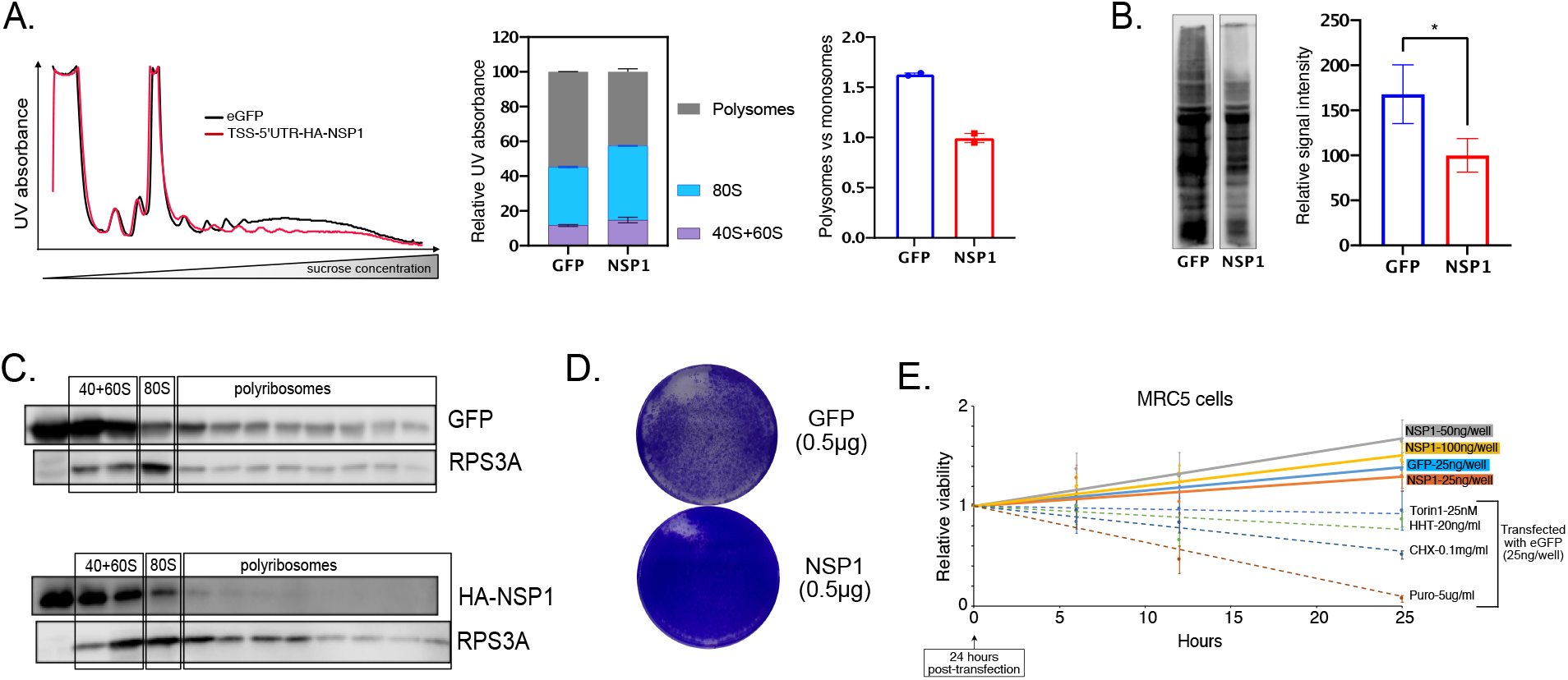
The impact of NSP1 on translation and survival. **A.** HEK293 cells were transfected with plasmids encoding for either HA-NSP1 or eGFP, as a control. Twenty four hours later, the cells were collected, lysed and subjected to polysomal profiling. The left panel displays continuous UV absorbance of both gradients of one out of two independent experiments. The middle panel shows the signal distribution between the different grouped fractions. The right panel directly compares the polysomes vs monosomes fractions. In all panels, bars are SD of n=2. See also Fig. S2C for analysis of MCF7 cells. **B.** HEK293 cells were transfected with plasmids encoding either eGFP or HA-NSP1. After 24 hours, puromycin (10μg/ml) was added for 5 mins, and the cells were collected on ice, lysed, and 50μg of total lysates were resolved on 10% SDS-PAGE and probed with anti-puromycin antibodies. The relative signals obtained from the different lanes were analyzed and plotted on the right panel; n=4, bar represents SD. See also Fig. S2A for the original images and S2B for expression of specific proteins. **C.** MCF7 cells were transfected as in B and after 24 hours lysed and subjected to polysomal isolation. The total proteins extracted from the collected polysomal fractions were separated on SDS-PAGE and probed to detect the indicated proteins. **D.** HEK293 cells were grown in 6-wells dishes, transfected with either eGFP or TSS-5’UTR-HA-NSP1 (0.5μg/well) and allowed to grow for additional 3 days, after which they were fixed and stained. See also Fig. S2D, E for more images. **E.** MRC5 cells were seeded in 96-well plates and transfected with the indicated amounts of plasmids encoding for either eGFP or HA-NSP1. Twenty-four hours post-transfection (t=0), the cells were subjected to proliferation assay at indicated time points. As positive controls, cells were treated with known inhibitors of translation at t=0 at indicated concentrations. N=3, bars represent SE. See also Fig. S2F for a similar analysis of Vero cells.

### Resistance of viral mRNAs to NSP1-mediated inhibition of translation is conferred by a precisely located RNA sequence motif

Next, we investigated the translational features of the viral 5’UTRs. Viral mRNAs produced in SARS-CoV-2-infected cells bear either long genomic (265-nt long) or shorter sub-genomic (app. 70-75-nt long) 5’UTRs, the latter being produced via discontinuous transcription (6). The genomic 5’UTR of SARS-CoV-2 precedes the two first open reading frames (ORF1ab) that cumulatively yield 16 non-structural proteins. The NSP1-encoding construct used in this study until now, encodes for a full genomic 5’UTR placed immediately after the transcription start site (TSS). Based on previous reports (13), we anticipated that replacing the native 5’UTR will expose NSP1 to self repression. Indeed, this manipulation significantly reduced the protein level of NSP1 (Fig. 3A), indicating that genomic 5’UTR of SARS-CoV-2 grants resistance to auto-inhibition. To test this further, we transfected HA-NSP1 into cells and 24h later transfected capped mRNAs encoding the Renilla luciferase (Rluc) under the control of the SARS-CoV-2 genomic, sub-genomic or a control (90 nt-long) 5’UTRs. While mRNA encoding the control 5’UTR was inhibited by NSP1, both the genomic and sub-genomic 5’UTRs not only conferred full protection but in fact boosted the reporter expression (Fig. 3B, blue columns). To understand how viral 5’UTRs escape the inhibition by NSP1, we focused on the first stem-loop structural element within the 5’UTR (SL1, Fig. 3C, left panel), which was previously reported to bind NSP1 of SARS-COV-1 and also suggested to play a role in SARS-CoV-2 (12, 13, 22). First, we compromised its structure in the context of sub-genomic (72-nt long) 5’UTR by introducing five point-mutations in the distal part of the stem (S1mut, Fig. 3C). Interestingly, disrupting this element did not interfere with the protection of this element from NSP1 inhibition and even slightly enhanced it (Fig. 3B, yellow columns), uncoupling the structural integrity from NSP1-mediated repression. Notably, further alteration of the sequence comprising SL1 to restore the stem-loop structure significantly disrupted the protective effect against NSP1 (S1_comp mRNA, Figs. 3C and B). These findings strongly indicate that the ability to oppose NSP1 inhibition does not rely on the secondary structure of the SL1 element, but rather on its primary RNA sequence. To check further if cap-proximal stem-loop structures may protect mRNAs from NSP1, we exchanged the CoV-2-derived SL1 by an even longer stem-loop element derived from the Chikungunia virus (Chikungunia SL1 mRNA, Figs. 3B,C). Indeed, this element was inefficient in protecting the reporter transcript, indicating that cap-proximal secondary structures are unlikely to relief NSP1-mediated inhibition. Importantly, insertion of 10 nt between the cap and the SL1 element reduced the expression of the reporter mRNA (72+10 mRNA, Figs. 3B, C), indicating the importance of the location of this element relative to the 5’end. Taken together, the results suggest that the sequence comprising the SL1 element and its location relative to the 5’cap are important features for resisting NSP1 inhibition.

**Figure 3:**
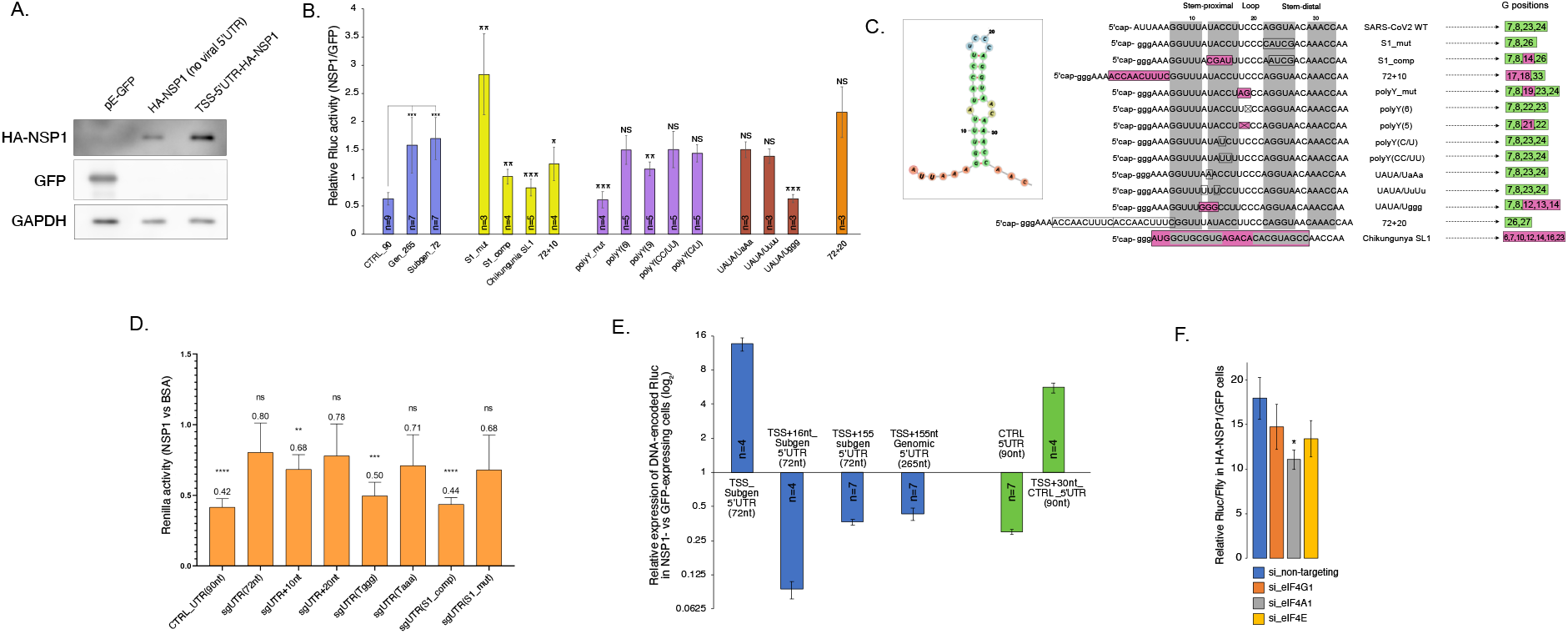
Identification of RNA sequence that protects from NSP1-mediated translational repression. **A.** MRC5 cells were transfected with plasmids encoding for eGFP, HA-NSP1 or HA-NSP1 with its native 5’UTR immediately following the TSS (TSS-5’UTR-HA-NSP1). After 24 hours, the cells collected and 50μg from the extracted total proteins were resolved on 9% SDS-PAGE and probed to detect the indicated proteins. **B.** Impact of NSP1 on the expression of reporter mRNAs. MRC5 cells were transfected with plasmids encoding either eGFP or TSS-5’UTR-HA-NSP1. After 20 hours, the indicated mRNA reporters bearing both 5’cap and poly(A) tail, were transfected and the cells were subjected to luciferase assay 7 hours later. Number of independent biological repeats is indicated for each reporter, bars indicate SE. p values indicated for the blue columns refer to the reporter bearing the control 5’UTR (CTRL_90), while for the rest of the columns it refers to the reporter encoding the sub-genomic SARS-CoV2 5’UTR (Subgen_72). **C.** The window on the left represents the SL1 element’s structure. The middle panel shows detailed schemes representing the manipulations done on the 5’cap-proximal sequence of the viral 5’UTR. Changed moieties are squared; colorless squares imply no change to expression, pink squares imply reduced expression, grey shades mark the nucleotides participating in stem structures. The rightmost panel shows the positions of guanosine moieties within the corresponding sequences relative to the 5’cap. Green color indicates unperturbed expression, while new locations of the “G” moieties that reduce expression are indicated in pink. **D.** *In-vitro* transcribed Rluc mRNA reporters bearing both 5’caps and poly(A) tails were added to RRL pre-incubated with either HA-NSP1 or BSA purified proteins. Rluc activity was tested after 60 minutes of incubation. The graph shows the ability of NSP1 to inhibit the different mRNA reporters at the translational level, n=3, bars represent SD. **E.** Plasmids encoding for the indicated configurations of 5’UTRs followed by the reporter Rluc gene were co-transfected with a plasmid encoding either for HA-NSP1 or eGFP into MRC5 cells; Rluc activity was tested after 48 hours following transfections. Number of biological repeats (n) is indicated separately for each combination; bars represent SE. **F.** MRC5 cells were transfected with the indicated siRNAs and after 48 hours transfected again with plasmids encoding i) either TSS-5’UTR-HA-NSP1 or eGFP, ii) TSS-5’UTR-Rluc and, iii) Firefly (Ffly). After additional 24 hours, luminescence was tested and Rluc values were normalized by Ffly signal and compared to cells transfected with the control non-targeting siRNAs for statistical analysis. N=3, bars represent SE.

To better understand the code necessary for mRNA protection, we focused on two elements that we identified in the sequence comprising the SL1 element: i) a stretch of seven consecutive pyrimidines located in the proximal part of the stem and the loop (nt. 15-21, Fig. 3C), and ii) a UAUA motif immediately preceding the polypyrimidine stretch (nt. 11-14). First, we examined the polypyrimidine stretch because its disruption (in the SL1_comp construct) greatly reduced the protection from NSP1. By manipulating this element, we found that: i) introduction of two purine (AG) residues in the middle of the stretch significantly reduced mRNA protection (see polyY_mut); ii) shortening the stretch from 7 to 6 and 5 pyrimidines gradually impaired the protection (see polyY(6) and polyY(5)); and, iii) the precise pyrimidine sequence at the beginning of the stretch had no effect (see polyY(CC/UU) and polyY(C/U)). These results indicated that the presence of polypyrimidine stretch and its length have a role in the NSP1 protection.

We next mutated the UAUA sequence to yield UAaA, UuUu or Uggg. Intriguingly, we found that while the first two manipulations had no significant effect, insertion of guanosines turned the mRNA extremely sensitive to NSP1-mediated repression (see UAUA/Uggg, Figs. 3B, C). Notably, in this mutant, the polypyrimidine stretch was intact, strongly indicating that it is insufficient to grant NSP1 resistance. Additionally, these results suggest that the presence of guanosines, but not adenosines or thymidines, in the region of SL1 may predispose mRNAs to NSP1-mediated repression. To test this further, we mapped the precise locations of guanosines in the different mRNA variants that we tested above (Fig. 3C, “G positions” column). While the original viral 5’UTR lacks guanosines between positions 9-22, all mutant 5’UTR versions that were repressed by NSP1 (e.g., S1_comp, WT+10, etc.), involved insertion of guanosines between these precise positions (magenta squares). Intriguingly, manipulations that did not alter the positions of guanosine residues (empty squares) rendered mRNAs protected to the levels comparable to the original 5’UTR. To test if the precise positioning of guanosines could play a role in NSP1 resistance, we duplicated the inserted 10 nucleotides in the WT+10 mRNA. This new version of the 5’UTR did not introduce any new sequences, but pushed the existing guanidines from positions 17, 18 to 26 and 27 relative to the 5’-cap. Strikingly, this repetitive addition of 10 nucleotides fully restored the NSP1 resistance (Fig. 3B, 72+20). Since these 10 nucleotides, which are guanosine-free, were already present in the WT+10 mRNA, this result indicates that this sequence *per se* is not sufficient to resist NSP1 inhibition but is important to create a guanosine-free stretch precisely located at positions 9-23 relative to the 5’cap. To test if 5’cap-proximal guanosine-free stretches are conserved in coronaviruses, we mapped G-free regions in the 5’UTRs of several alpha-and beta-coronaviruses (Fig. S3A). Indeed, we found that multiple CoV-derived 5’UTRs encode for cap-proximal guanosine-free stretches, indicating the conservation and, therefore, importance of this feature. Importantly, we recapitulated this phenomenon using rabbit reticulocyte lysate *in vitro* translation system (Figs. 3D), indicating that this regulation occurs at the level of translation. We also confirmed that guanosine residues located between the positions 11-20 relative to the 5’cap structure do not alter mRNA stability in the presence of NSP1 (Fig. S3B).

To test these conclusions further, we decided to apply them on plasmid-encoded genes. To our knowledge, NSP1 efficiently represses expression of DNA-encoded constructs (e.g, Fig. 1D). According to our findings, we constructed a DNA-encoded reporter gene preceded by the sub-genomic version of the viral 5’UTR immediately downstream the TSS in order to preserve the guanosine-less state at the beginning of the transcript and examined its expression in the presence of NSP1. Indeed, we found that the precise location of the viral 5’UTR sequence not only fully protected the gene from NSP1 repression, but in fact enhanced its expression by ∼13-fold in NSP1-expressing cells (Fig. 3E, TSS_subgen 5’UTR). Moving the viral 5’UTRs away from the TSS by introducing either 16 or 155 bases strongly reduced the expression of the respective DNA-encoded reporter genes and promoted vulnerability to NSP1 inhibition, both in the context of genomic and sub-genomic 5’UTRs (Fig. 3E, blue columns). These results strongly support the notion that the precise location of the viral 5’UTR relative to the cap structure is crucial to escape NSP1-mediated repression. To examine whether the first 30 nt of the viral 5’UTR that encode for guanosine-free stretch of nucleotides are sufficient for this effect, we inserted them after the TSS of the control 5’UTR, which is otherwise NSP1-sensitive (Fig. 3E, green columns). This manipulation not only abolished its inhibition by NSP1, but also boosted it’s expression ∼5-fold (Fig. 3E, green columns). While this level of induction was somewhat lower compared to the native sub-genomic SARS-CoV-2 5’UTR (∼15-fold induction), it nevertheless supported the idea that a guanosine-free RNA sequence precisely located relative to the 5’end is sufficient to confer NSP1 resistance.

Lastly, we assumed that successful translation in NSP1-expressing cells may require unwinding of the secondary structure of SL1 in order to expose its primary RNA sequence. We therefore knocked-down the different elements of the eIF4F complex and tested the effect on NSP1-mediated boost of the DNA construct encoding the sub-genomic 5’UTR, which is efficiently translated in NSP1-expressed cells (Fig. 3E, TSS_subgen 5’UTR). Interestingly, we found that repression of the RNA helicase component, eIF4A, had the strongest negative effect on the ability of NSP1 to promote the expression of this gene (Fig. 3F), supporting the notion of the necessity to unwind the secondary SL1 structure in order to enable efficient protection from NSP1 inhibition. All together, these results suggest that protection of mRNAs from NSP1-mediated inhibition of translation is mediated by a precisely located cap-proximal guanosine-deficient RNA sequences.

### The genomic 5’UTR mediates 5’cap-independent translation initiation

Next, we investigated the translational properties of the genomic (265-nt) and sub-genomic (72-nt) 5’UTRs of SARS-CoV-2. We reasoned that since the cap-proximal sequence that grants protection from NSP1-mediated repression is present in both versions, the longer genomic version may encode for additional translational features. Particularly, we tested its capacity to promote cap-independent translation initiation, which is a feature common to multiple positive single-stranded RNA viruses. To this end, we subcloned the genomic 5’UTR into a bi-cistronic system (Fig. 4A) and tested its cap-independent activity in MCF7 cells. Interestingly, the presence of the genomic 5’UTR markedly enhanced the expression of the downstream gene, in a manner comparable to the well-characterized EMCV-derived IRES element (Fig.4B, left, blue columns). Importantly, we observed a similar effect in MRC5 cells (Fig.4B, right panel, blue columns), suggesting that this feature is not cell-type specific. Neither element showed activity when cloned in the opposite configuration (i.e., flipped), indicating the importance of the primary RNA sequences for both elements. Characteristically to the known IRES elements, treatment with Torin1, a pharmacological agent that reduces availability of eIF4E and inhibits cap-dependent translation, potentiated the activity of both elements (Fig. 4B, orange columns in both graphs), indicating their ability to sustain translation even upon inhibition of cap-dependent initiation. To directly test the ability of the genomic 5’UTR to drive cap-independent translation, we transfected uncapped reporter mRNAs bearing different viral 5’UTRs into cells and assayed the protein activity of the corresponding genes. Indeed, we observed a relatively enhanced (app. 3.5-fold) expression of the reporter mRNA bearing the genomic 5’UTR version (Fig. 4C). This feature disappeared when the sequence was cloned in its anti-sense configuration (Fig. S4A), indicating sequence specificity. To further validate these findings, we tested the sensitivity of viral 5’UTRs to knock-down of eIF4F components (Fig. S4B). Knock-down of either eIF4A1 or eIF4G1 reduced the translatability of all reporters, whereas knock-down of the cap-binding eIF4E subunit had a differential effect on viral 5’UTRs (Fig. 4D). eIF4E deficiency dramatically reduced the expression of mRNA encoding the sub-genomic 5’UTR while the genomic 5’UTR exhibited relative insensitivity, confirming its ability to resist attenuation of cap-dependent translation.

**Figure 4:**
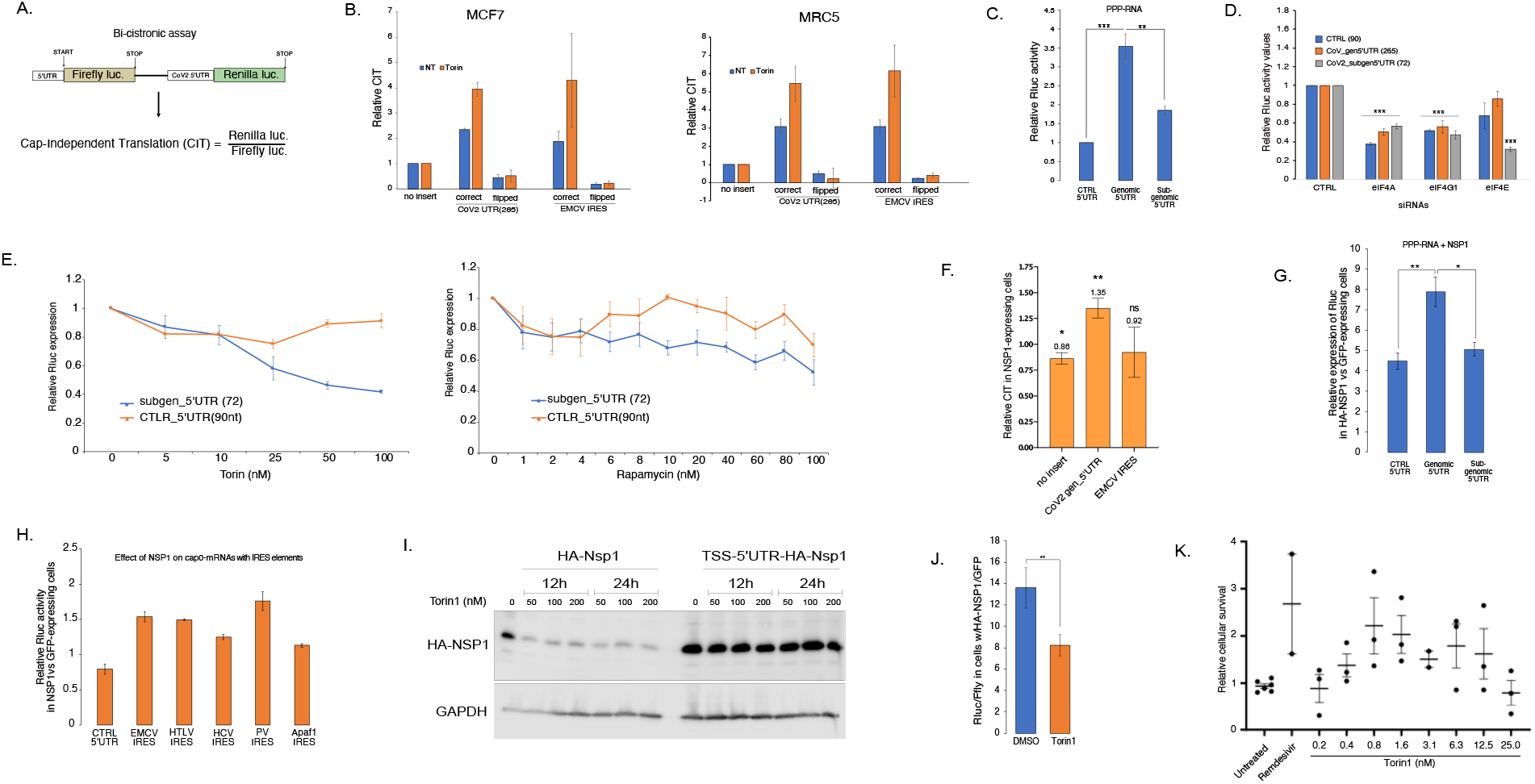
The cap-independent translational features of SARS-CoV-2 5’UTRs. **A.** Schematic representation of the bi-cistronic assay and the calculation of cap-independent translation. **B.** MCF7 (left panel) or MRC5 (right panel) cells were transfected with bi-cistronic assay plasmids encoding for the SARS-CoV-2 genomic 5’UTR or EMCV-derived IRES element as positive control. Both elements were introduced in their correct or flipped (i.e. anti-sense) configuration as a control for non-specific activity. Cells were grown for 2 days in normal medium (NT) or supplied with Torin1 (10nM), after which Rluc and Ffly activities were assayed; n=3, bars represent SE. **C.** MRC5 cells were transfected with the indicated uncapped (PPP) in-vitro transcribed mRNAs bearing poly(A) tails. Rluc activity was tested 5 hours after the transfection and presented relatively to the control plasmid derived 5’UTR (90nt). N=4, bars represent SE. **D.** MRC5 cells were transfected with the indicated siRNAs and, after 65 hours, with *in-vitro* transcribed mRNA reporters bearing both 5’caps and poly(A) tails. Rluc activity was tested 7 hours after the second transfection and presented relatively to the cells expressing the control siRNA; n=4, bars represent SE. **E.** MRC5 cells were transfected with the indicated *in-vitro* transcribed mRNA reporters bearing both 5’caps and poly(A) tails. After 2 hours, Torin1 (right panel) or Rapamycin (left panel) were added to yield the indicated final concentrations and Rluc activity was tested after 5 hours; n=3, bars represent SE. **F.** MRC5 cells were transfected with i) the indicated bi-cistronic plasmids and ii) plasmid encoding either HA-NSP1 or eGFP. Luminescence was assayed 24 hours later, n=3, bars represent SD. **G.** MRC5 cells were transfected with plasmids encoding either HA-NSP1 or eGFP. After 24 hours, uncapped RNAs with poly(A) tails encoding for indicated 5’UTRs and a downstream Rluc gene were transfected and Rluc activity was assayed 7 hours later; n=4, bars represent SE. **H.** MRC5 cells were transfected with plasmids encoding either HA-NSP1 or eGFP. After 24 hours, mRNAs with cap0 structures and poly(A) tails encoding for indicated IRES elements and a downstream Rluc gene were transfected and Rluc activity was assayed 7 hours later. N=2, bars represent SE. **I.** HEK293 cells were transfected with plasmids encoding HA-NSP1 either without viral 5’UTR or with its native genomic 5’UTR introduced immediately following the transcription start site. The cells were grown for 40 hours post-transfection, out of which Torin1 was added during the last 12 or 24 hours at the indicated concentrations. After harvest, total lysates were separated on 9% SDS-PAGE and probed with anti-HA and anti-GAPDH antibodies. **J.** MRC5 cells were transfected with the mix of plasmids encoding i) either TSS-5’UTR-HA-NSP1 or eGFP, ii) TSS-5’UTR-Rluc and, iii) Firefly (Ffly). After 4 hours of transfection, the medium was replaced and either DMSO or Torin1 (25nM) were added. The cells were collected after 24 hours, lysed and subjected to luminescence analysis. Rluc values were normalized against Ffly signal, n=3, bars represent SE. **K.** Vero E6 cells were treated with the indicated final concentrations of Torin1 for 1 hour, infected with SARS-CoV-2 at MOI=0.01-0.015 and incubated for three days in the presence of Torin1. Following this time, the relative cytopathic effect of the virus was tested. As a positive control, cells were treated with Remdesivir (0.3mM), for negative relative control cells were not treated prior to infection; n=2 or 3, bars indicate SE.

To further examine the sensitivity of the sub-genomic version of the viral 5’UTR to inhibition of cap-dependent translation, we applied two mTOR inhibitors, Torin1 and Rapamycin, and elucidated their effect on the translation of the reporter gene driven by the sub-genomic 5’UTR. Indeed, this 5’UTR was remarkably vulnerable to both inhibitors (Fig. 4E), confirming that translation of transcripts with sub-genomic 5’UTRs heavily relies on cap-dependent initiation. Overall, these results suggest that while the genomic 5’UTR of SARS-CoV-2 has the capacity to promote IRES-like cap-independent translation initiation, its sub-genomic shorter version exhibits a particular dependency on eIF4E availability.

### NSP1 enhances cap-independent translation

Despite that cap-independent translation is relatively inefficient compared to the cap-dependent mode, it is frequently employed by viruses in order to counteract the inhibition of cap-dependent translation, which is a part of anti-viral cellular defence (23). To test the effect of NSP1 on cap-independent translation, we employed the bi-cistronic assay and found that NSP1 moderately enhanced the cap-independent translation directed by the genomic 5’UTR (Fig. 4F). Remarkably, testing the effect of NSP1 on uncapped mRNAs, we found that it significantly boosted the expression of all tested transcripts, even those not encoding for viral 5’UTRs (Fig. 4G). The differences between the degree of activation in these experiments may stem from the ability of NSP1 to repress gene expression by multiple pathways, such as nuclear export (24). The latter result strongly indicates that NSP1 non-specifically enhances cap-independent translation. To test this assumption further, we co-transfected HA-NSP1-expressing cells with capped mRNAs encoding for different IRES elements in their 5’UTRs. Indeed, we found that although neither of the constructs encoded for cap-proximal guanosine-deficient elements that could protect mRNAs from NSP1-mediated repression, all tested IRES elements, including the mammalian Apaf1-derived, enhanced the expression of their corresponding transcripts in the presence of NSP1 (Fig. 4H). These results confirm the observation that a mere expression of NSP1 may generally enhance cap-independent translation.

Our findings strongly suggest that expression of NSP1 itself, which is preceded by the genomic 5’UTR, may have two important protective features. First, the cap-proximal guanosine-deficient element protects all viral transcripts from NSP1-mediated inhibition. Second, the IRES-like activity of the genomic 5’ UTR enables expression upon inhibition of cap-dependent translation. To examine the extent by which these features affect NSP1 expression, as a proxy of viral genes encoded by ORF1ab, we employed two versions of NSP1: one bearing the native genomic 5’UTR correctly positioned relative to the 5’cap and the second under a control 5’UTR. Upon transfection into cells and application of Torin1 we found that in contrast to the control 5’UTR, the native 5’UTR not only enhanced NSP1 expression, but also retained its high levels despite treatment with Torin-1 (Fig. 4I).

Considering the efficient expression of NSP1 under conditions of inhibited cap-dependent translation, we examined the impact of NSP1 on the expression of reporter mRNAs driven by the sub-genomic 5’UTRs, which strongly depends on eIF4E (Figs. 4D, E), upon Torin1 treatment. Interestingly, we found that although the ability of NSP1 to promote the expression of sub-genomic 5’UTR-encoding transcripts was lower relative to untreated cells, it was still potent and boosted the expression by 8-fold (Fig. 4J).

Considering the negative impact of Torin1 on the expression of sub-genomic 5’UTRs and the ability of NSP1 to counteract this inhibition, we tested the effect of Torin1 treatment on virus propagation in cells. For this purpose, we treated Vero E6 cells with different concentrations of Torin1, infected them with SARS-CoV-2 and monitored the cytotoxic effect of the virus after 72 hours. Importantly, we found that mTOR inhibition enhanced the survival of the infected cells at relatively low doses (i.e., 0.8-12.5 nM), suggesting that mTOR inhibition may have an anti-viral effect, while at higher concentrations it was toxic for the cells (Fig. 4K).

## Discussion

NSP1 protein encoded by alpha- and beta-coronaviruses plays a central role in the virus infectivity in general and the ability to control the host translation machinery in particular (25). By binding the mRNA entry channel of the small ribosomal subunit, NSP1 both hinders the translation of cellular mRNAs and induces their degradation by a still unclear mechanism. In this study we elucidated the molecular features that enable SARS-CoV-2 to hijack host translation to prevent efficient expression of host genes and, at the same time, to enable its own mRNAs to avoid these inhibitory mechanisms. We found that a mere expression of NSP1 is sufficient to target the vast majority of the host mRNAs for degradation, consistent with recent studies (9–11). Our findings further reveal that NSP1-induced degradation is specific to spliced mRNAs, rendering single-exon mRNAs relatively stable. Interestingly, interferon alpha and beta genes, that comprise the first line of the host anti-viral response, are all intron-less and therefore are likely to be resilient to NSP1-induced mRNA degradation. To further limit the expression of the host genes, SARS-CoV-2 has evolved an effective suppression strategy at the translation level, which hampers the entrance of mRNAs into the ribosome (7, 8, 21). We demonstrate that expression of NSP1 does not completely shut down the host translation, as evident from the polysomal profiles and puromycin labeling experiments (Figs. 2A, B) but rather attenuates the translation initiation step (21, 26). In support of that, we found that NSP1 is enriched in the initiating fractions (Fig. 2C) leading to enhanced RNA content in fractions representing 80S (Figs. 2A and S2C). These findings are in agreement with recent reports of relatively unperturbed translation elongation dynamics in both infected cells (10) and upon NSP1 expression (11). Moreover, recent studies demonstrated that NSP1 may profoundly interfere with the nuclear-cytoplasmic shuttling (9, 24, 27). Surprisingly, despite these prominent effects on host gene expression, we detected no cytotoxic effects of NSP1 expression in multiple tested cell types (Figs. 2D and S2D-F). While the precise roots of this phenomenon remain to be understood, it is likely that the lack of toxicity of NSP1 is important to ensure successful viral propagation.

Overall, we show that SARS-CoV-2 employs at least three strategies to promote efficient translation of its own mRNAs. First, by inducing degradation of most cellular mRNAs (Fig. 1C), NSP1 reduces the competition of the viral intron-less transcripts with cellular mRNAs for accessing the ribosomes. Second, we identified a regulatory sequence in SARS-CoV-2 5’UTRs that is sufficient to confer protection from NSP1-mediated inhibition of translation. A detailed characterization of this element revealed that it overlaps the SL1 element, as recently reported (13), but our results suggest that the primary sequence mediates the protective effect, rather than the secondary structure that it creates. This finding is supported by the strong dependency of viral transcripts on eIF4A (Fig. 3F and (28, 29) that might be needed to unwind the SL1 structure and expose its sequence. We show that the G-less nature of this motif and its proximity to the 5’cap structure are the most important features that protect viral mRNAs from NSP1-mediated repression, and these features are conserved among several coronaviruses (Fig. S3A). Third, we found an IRES-like activity of the genomic viral 5’UTR that bypasses reduced eIF4E availability (Fig. 4B). This feature enables cap-independent translation of the first two viral ORFs that encode 16 non-structural viral proteins. Importantly, the sub-genomic versions of the viral 5’UTR are highly eIF4E dependent (Fig. 4E), but efficient cap-independent expression of NSP1 alleviates this dependency to some extent (Fig. 4J). This phenomenon may contribute to the sensitivity of SARS-CoV-2 to eIF4E inhibitors, at least in culture (Fig. 4K). Notably, the efficacy of mTOR targeting as a potential anti-COVID19 treatment is currently under debate (30–33), and our findings support the possible efficacy of such therapy (Fig. 4K).

The ability of the genomic 5’UTR of SARS-CoV-2 to mediate cap-independent translation initiation is rather intriguing, particularly because all viral transcripts bear 5’cap structures. This feature might be necessary at the initial stages of infection, which provoke interferon response, leading to inhibition of cap-dependent translation (1, 23). Under these conditions, the genomic 5’UTR may act in an IRES-like manner, promoting cap-independent translation of ORF1ab and synthesis of multiple non-structural proteins necessary to counteract the host defense (34) and exploit the cellular machineries. Interestingly, we found that mere expression of NSP1 promotes cap-independent translation, enabling relatively higher translational yields from both uncapped and IRES-bearing mRNAs (Fig. 4G,H). These findings corroborate the notion that SARS-CoV-2 may be well prepared for hijacking the host translational machinery under conditions of inhibited cap-dependent translation. However, at the later stages of infection, cap-dependent initiation is necessary to translate viral mRNAs bearing short sub-genomic 5’UTRs, which we found to strongly depend upon eIF4E availability. This might be an Achilles heel of viral replication and the reason for the sensitivity of SARS-CoV-2 to pharmacological mTOR inhibitors (33). Moreover, this study highlights the roles of NSP1 and the viral 5’UTR-located protective element in the translation regulation exerted by the virus. These features may be further exploited as major vulnerabilities of SARS-CoV-2 and are, therefore, excellent targets for general therapeutic applications against coronaviruses aiming to impair the early essential stages of viral gene expression and facilitate efficient anti-viral response by the host.

## Data availability

All data regarding the genome-wide determination of mRNA stability presented in Figure 1 will be made public upon acceptance of the manuscript.

## Funding

This work was supported by grants from the Israel Science Foundation KillCorona fund (3694/20) the Israel Science Foundation (#843/17); the Minerva Foundation (#713877) and by Weizmann Institute internal grants from CoronaVirus Fund; Estate of Albert Engleman; Estate of David Levinson. R.D. is the incumbent of the Ruth and Leonard Simon Chair of Cancer Research.

## Materials and Methods

### Reagents

#### DNA constructs

To create HA-tagged NSP1 lacking viral 5’UTR for expression in cell culture (pCRUZ-HA-NSP1), NSP1 was obtained from the Forchheimer plasmid bank (WIS, Israel) and cloned into pCRUZ-HA (sc-5045) plasmid (Santa Cruz Biotechnology) using restriction-free cloning and primers #1,2 (here and on, see Table S1 for primers’ sequences). To create His-tagged NSP1 lacking viral 5’UTR for bacterial expression, NSP1 was PCR amplified using primers #3,4 and inserted into pET28 plasmid with 14×His-bdSumo tag (14). The insertion of NSP1 removed the bdSumo but maintained the 14×His tag. To create HA-tagged NSP1 with the viral 5’UTR for expression in cell culture (pcDNA3.1-TSS-5’UTR-HA-NSP1), HA-NSP1 was PCR-amplified from the pCRUZ-HA-NSP1 plasmid using primers #5,6 and introduced immediately after the transcription start site of the pcDNA3.1(-) plasmid using restriction-free cloning. This plasmid (pcDNA3.1-TSS-5’UTR-HA-NSP1) was used in this study to express HA-NSP1 in mammalian cultured cells, unless indicated otherwise. The genomic 5’UTR was created by sequential annealing and amplification of primers #13-17. The combined 5’UTR was introduced between KpnI and BamHI sites of pcDNA5/FRT/TO plasmid; Renilla (Rluc) reporter gene was introduced into the same plasmid using BamHI-NotI sites. For bi-cistronic plasmids, Firefly (Ffly) reporter gene bearing Hisx6 tag was amplified using primers #18,19 and cloned between HindIII-KpnI sites of pcDNA5/FRT/TO plasmid. This plasmid was used to both express Firefly gene in cells and synthesize Firefly mRNA *in vitro* using T7 promoter. Rluc was amplified using primers #20,21 and introduced using XhoI site of the same plasmid to create a bi-cistronic expression vector where both reporters have distinct translation start and stop codons. After cloning, the XhoI site was preserved only before the Rluc gene and used for insertion of tested DNA sequences in both orientations. For this purpose, the genomic 5’UTR was amplified using primers #22,23, EMCV IRES with #24,25. To create a plasmid with sub-genomic 5’UTR fused to Rluc, the Rluc gene was first amplified with primers #26,27 and the resulting product was further re-amplified with primers # 27,28. The resulting product was used to run a PCR using pcDNA5/FRT/TO plasmid as a template. The resulting plasmid had 16nt between the transcription start site (TSS) and the beginning of the viral 5’UTR. To create plasmid bearing 155nt between the TSS and the viral 5’UTR, the cassette including the sub-genomic 5’UTR and Rluc gene was amplified using primers #13,29 and inserted between KpnI-NotI sites of pcDNA5/FRT/TO. The same strategy was used to create plasmid bearing the genomic 5’UTR. To create a plasmid with viral 5’UTR placed immediately after the TSS, primers # 13,30 were used to amplify the 5’UTR-Rluc cassette and the product was re-amplified using primers #31,32. The resulting product was used in a PCR to amplify pcDNA3.1(-) plasmid as template. All constructed plasmids were subjected to partial Sanger sequencing to ensure correctness of the sequences.

#### Restriction-free cloning

For restriction-free cloning, a PCR product with ends overlapping the destination plasmid was used for a subsequent PCR using the destination plasmid as a template. The resulting product was cleaned, concentrated, treated with DpnI enzyme for 1-2 hours and transformed into bacteria. Grown colonies were screened using colony PCR using HyTaq mix (HyLabs, Israel).

#### Plasmid DNA preparation

Plasmid DNA of interest was transformed into competent DH5 alpha bacteria using heat shock method and plated on LB plates supplemented with either Ampicillin (200μg/ml) or Kanamycin (50μg/ml). Grown colonies were isolated, grown overnight in liquid LB medium supplemented with the relevant antibiotic, and plasmid DNA was extracted using either miniprep or maxiprep kits.

#### RNA constructs

DNA templates bearing T7 promoter were produced by PCR amplification (Kappa HiFi Hotstart, Roche) of plasmids encoding for Renilla reporter using primers detailed in Table S1. Typically, primer #33 was used to add poly(A) tail of 30nt and primers #34-47 introduced both T7 promoter and the indicated sequence manipulations. PCR products were cleaned using columns (Qiagen) and used for *in-vitro* transcription reactions, which were done using RiboMAX kit (Promega) according to the manufacturer’s instructions. The reactions were treated with DNAse (15 min at 37°C) and remaining RNA was recovered using Direct-zol RNA mini prep kit (Zymo research) according to the protocol. When needed, RNA was capped Vaccinia capping kit (NEB) and recovered using Directzol RNA mini prep kit (Zymo research).

#### RNA isolation and qRT-PCR assay

RNA was isolated from cultured cells using BIO TRI RNA reagent (Bio-lab) and Direct-zol RNA MiniPrep kit (Zymo research), as instructed by the manufacturer. RNA was typically reconstituted in 25μl of sterile nuclease-free water (Bio-lab) and stored at −20°C. Reverse transcription was done using High Capacity cDNA Reverse Transcription Kit (Applied Biosystem), according to the manual. Typically, 5μl of total RNA were taken for a single RT reaction and diluted 1:5 with water afterwards. Quantitative real time PCR (qRT-PCR) experiments were performed in total reaction volume of 10μl with qPCRBIO SyGreen Blue Mix Hi-ROX (PCR BIOSYSTEMS) reagent on 384-wells plates (Axygen) using Viia7 (Thermo Fisher Scientific) instrument in standard conditions.

#### Nsp1 protein purification for in vitro assay

The pET28 plasmid encoding for His-tagged NSP1 was transformed by heat shock into BL-21 E.coli and a single grown colony was inoculated into 5ml of LB supplemented with Kanamycin (50μg/ml) and incubated shaking at 37°C. After 8 hours, the bacteria were transferred into Erlenmeyer flask with 1 liter of LB supplemented with Kanamycin and grown to O.D ∼0.5 at 37°C. IPTG (0.5 mM) was added to the medium and incubated orbitally shaking at 37°C overnight. The bacteria were pelleted (4000RPM, 20 minutes) and resuspended in 20ml of resuspension buffer (500mM NaCl, 30mM Hepes, 5mM MgCl). The resuspended bacteria were sonicated (probe sonicator, 12 cycles of 30 sec on + 30 sec off) and centrifuged to pellet and remove the cellular debris. Supernatant was then loaded on 500uL Nickel-NTA beads, washed 4 times with 10ml wash buffer (resuspension buffer with 20mM imidazole) and eluted with 5 ml resuspension buffer supplemented with 300mM imidazole. The eluate was subjected to buffer exchange in a dialysis bag (10 KDa) against resuspension buffer without imidazole and used for *in vitro* assays. This protocol is based on previously reported procedure for recombinant NSP1 isolation (8).

#### In-vitro translation assay

*In vitro* translation in RNase-treated Rabbit Reticulocyte Lysate System (RRL, Promega) was carried out as recommended by the manufacturer with slight modifications. The final volume of the reaction mixtures was 12.5 μl. The mixtures were preincubated for 10 mins with either NSP1 or BSA protein (5 ng/μl). Indicated *in-vitro* transcribed and capped mRNAs were added to the RRL reactions (6.25 ng/μl) and incubated at 30 °C for 1 hour. Samples of 3 μl were then subjected to Renilla activity measurements.

#### mRNA library preparation for MARS-seq

Total RNA was isolated from HEK293T cells using Bio Tri RNA reagent (Bio-lab) and mRNA was captured using Oligo d(T)25 magnetic Beads (NEB) according to manufacturer’s protocol. Poly(A)-purified RNAs were taken for library preparation using a derivation of MARS-seq as described (15). Briefly, 10 ng of poly(A)+ RNA was taken for the first reverse transcription reaction using illumina barcoded RT1 primer. Resultant barcoded cDNA samples were subsequently pooled according to Ct values of house-keeping gene (GAPDH) (Quality control 1). Pooled cDNA was treated with Exonuclease I to remove excess primers followed by second strand synthesis using NEB SSS module enzyme mix. After that, *in-vitro* transcription was performed using NEB T7 RNA Pol mix to generate RNA, which was later fragmented and ligated to an adaptor consisting of RD2 using T4 RNA ligase I (NEB) followed by the second reverse transcription reaction. The library so formed was amplified using Kapa Hifi ready mix (Roche). The amplified RNA libraries were sequenced using high-throughput 75 bp kit (Illumina FC404-2005) on NEXTseq 500 sequencer.

#### Polysomal isolation

Cultured cells were treated with 100 μg/ml cycloheximide for 5 minutes and washed with cold polysome buffer (20mM Tris pH8, 140 mM KCl, 5 mM MgCl2 and 100 μg/ml cycloheximide). Cells were collected in 500-μl polysome buffer supplemented with 0.5% Triton, 0.5% DOC, 1.5 mM DTT, 150 units RNase inhibitor and 5-μl protease inhibitor cocktail. After mechanical disruption, the samples were centrifuged at 12000 rpm for 5 minutes at 4°C. The cleared lysates were loaded onto sucrose density gradient (10-50%) and centrifuged at 38000 rpm for 105 minutes at 4°C. Gradients were fractionated with continuous absorbance 254nm (A254) measurement using ISCO absorbance detector UA-6. Fractions were pooled according to their absorbance into free, light and heavy ribosomal fractions. A whole sum of absorbance (A254) for 80S peak was calculated that marked as monosomal fraction. Similarly, the total sum of absorbance (A254) for both light and heavy fractions was calculated as polysomal fraction. For protein extraction, polysomal fractions collected from sucrose gradients were treated with 0.25 volumes of ice-cold 100% Trichloroacetic acid and 0.05% sodium deoxycholate to precipitate total proteins for 30 mins on ice. The samples were then centrifuged at 4°C 20000 *g* for 30 mins. The protein pellets were carefully washed twice with pure acetone, air dried and reconstituted in a 2X sample loading buffer. The protein samples were loaded into 12% SDS-PAGE gel and subjected to Western blot analysis.

#### Puromycin labelling

Cells were transfected with plasmids encoding either HA-NSP1 or eGFP and 24 hours later, puromycin (10 μg/ml) was added for 5 mins. The cells were then collected on ice and lysed using RIPA buffer (Sigma) and subjected to 10% SDS-PAGE followed by Western blotting.

#### Western blot

Protein extracts were resolved on SDS-PAGE using standard equipment (Bio-Rad) and transferred to PVDF membranes in buffer containing 20% methanol. Quality of transfer was examined by Ponceau staining (0.1% ponceau in 20% acetic acid) and membranes were blocked in skim milk (5% w/v) solution. The membranes were probed with the indicated antibodies overnight at 4°C, washed in washing buffer and probed with secondary HRP-conjugated antibodies (Jackson Immunoresearch). Primary antibodies used: anti-puromycin antibodies (Millipore Cat#MABE343), anti-alpha-tubulin (Sigma Cat# T5168), anti-GFP (Abcam Cat#ab1218), anti-HA (Abcam Cat#ab9110), anti-RPS3A (Antobody Verify, Cat#AAS38561C). After treatment with ECL reagent (Azure Biosystems), images were captured using Licor Fc imaging system and the signal intensities were calculated using ImageStudio software.

#### Luminescence assay

The cells were lysed in reporter lysis buffer (Promega), according to the volumes recommended by the manufacturer. For Renilla substrate, CTZ reagent (Bio Gold) was dissolved in methanol to stock concentration of 2.5mg/ml and equilibrated with 0.1N HCl. It was further diluted 1:1000 in phosphate buffer containing 80 mM K2HPO4 and 20 mM KH2PO4. Signals were detected in white 96-well plates using Modulus microplate luminometer reader (Turner Biosystems) combining 5μl cell lysate and 50μl of CTZ solution.

### Biological resources

#### Cultured cell lines

MCF7 and HEK293 cells were from ATCC, MRC5 and Vero E6 cells were generously provided by Zvi Livneh and Yosef Shaul, respectively (WIS, Israel). The cells were grown in DMEM media (Gibco) supplemented with 10% bovine serum and 1% penicillin/streptomycin (Gibco) in 5%CO_2_-buffered incubators at 37°C. Cells were split twice per week and kept in culture for up to 8 weeks.

#### Cell viability upon SARS-CoV-2 infection

SARS-CoV-2 (GISAID accession EPI_ISL_406862) was kindly provided by Bundeswehr Institute of Microbiology, Munich, Germany. Virus stocks were propagated (four passages) and titered on Vero E6 cells (Vero E6, ATCC® CRL-1586™). Handling and working with SARS-CoV-2 virus were conducted in a BSL3 facility in accordance with the biosafety guidelines of the Israel Institute for Biological Research (IIBR). Vero E6 cells were seeded at a density of 3×10^4^ cells per well in 96-well plates. After overnight incubation, cells were treated in 3 replicates with Torin-1. Cells were infected 1 hour later with SARS-CoV-2 (MOI, 0.01 - 0.015). Cell viability was determined 72 hours after infection by using the Cell Proliferation Kit (XTT based, Biological Industries, Israel) according to manufacturer’s protocol. As a positive control, cells were treated with Remdesivir, for negative relative control cells were not treated prior to infection.

#### Cell growth kinetics and proliferation assays

To visualize cellular growth, cells were seeded in 6-wells plates, treated as indicated, and grown for 48-72 hours in optimal conditions. The cells were washed once in PBS, treated with PBS-formaldehyde (4%) for 20 mins at room temperature and stained with crystal violet stain (Merck). After extensive washing, the plates were air-dried and documented. To monitor growth kinetics, the cells were seeded on 96-well plates, treated as indicated and subjected to CellTiter-Glo luminescent cell viability assay (Promega) according to the manufacturer’s instructions.

#### Transfections

DNA plasmids were transfected using JetPrime reagent (Polyplus transfection). Typically, the medium was changed after 4 hours, and cells were grown for 24-72 hours prior to further analysis. siRNA transfections were done using Lipofectamine RNAiMax reagent (Thermofisher scientific). The media were replaced 24 hours after transfections and the cells were collected after additional 48 hours. When indicated, cells were manipulated (e.g., transfected with DNA plasmids) within this time. *In-vitro* synthesized mRNAs were transfected using Lipofectamine 2000 reagent (Thermofisher scientific) according to the protocol provided by the manufacturer. Briefly, mRNA was mixed with Opti-MEM solution (Thermofisher scientific) and Lipofectamine reagent and incubated at room temperature for 10 mins. The growing media were replaced with pre-warmed Opti-MEM solution and the transfection solution was added to cells. The transfection solution was replaced after two hours to pre-warmed growth media and the cells were incubated at optimal conditions for 5 hours prior to lysis and subsequent luminescence assay.

### Computational resources

Raw data were processed using UTAP (16) with default parameters. Corrected counts were normalized by mouse PolyA+ enriched RNA, which was mapped to mouse genome using STAR (17) not allowing mismatches. Percent of uniquely mapped reads per sample was used as a normalization factor. Biological replicates were averaged and means were used for fitting a nonlinear least-squares model assuming first-order decay kinetics: C=C0*e-kdecayt, while corrected counts at t=0 were used as C0 and inverse of standard errors of the corrected count means were used as weights for fitting. Half lives of all genes were then calculated using the following equation: t1/2=ln(2)/kdecay. Genes with mean corrected read count < 5 at t=0 were removed from subsequent analysis. Negative half life values and half lives > 24h were set to 24h.

### Statistical analysis

Statistical significance was calculated using student’s t-tests with one-tailed distribution. Significance symbols in all experiments are: *=p<0.05; **=p<0.01; ***=p<0.001.

## Notes

### Competing Interest Statement

The authors have declared no competing interest.

