## Supplemental Figures for "Cap-independent translation and a precisely localized RNA sequence enable SARS-CoV-2 to control host translation and escape anti-viral response"

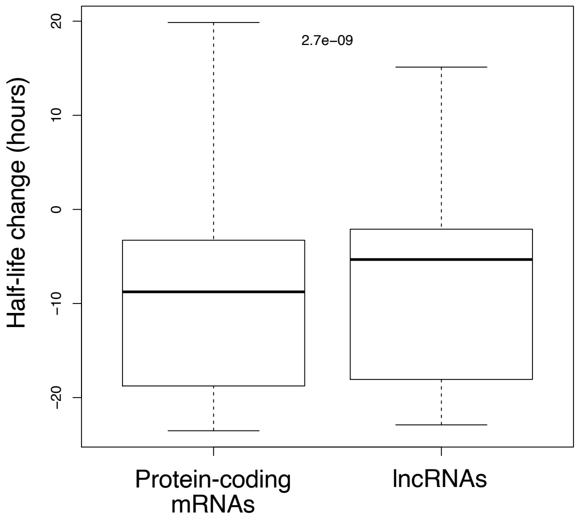


Figure S1: Inhibitory effects of NSP1. HEK293 cells were transfected and treated with ActD as detailed in Fig. 1B. The plot shows the negative effect of NSP1 on the stabilities of protein-coding versus long non-coding RNAs; n=2.

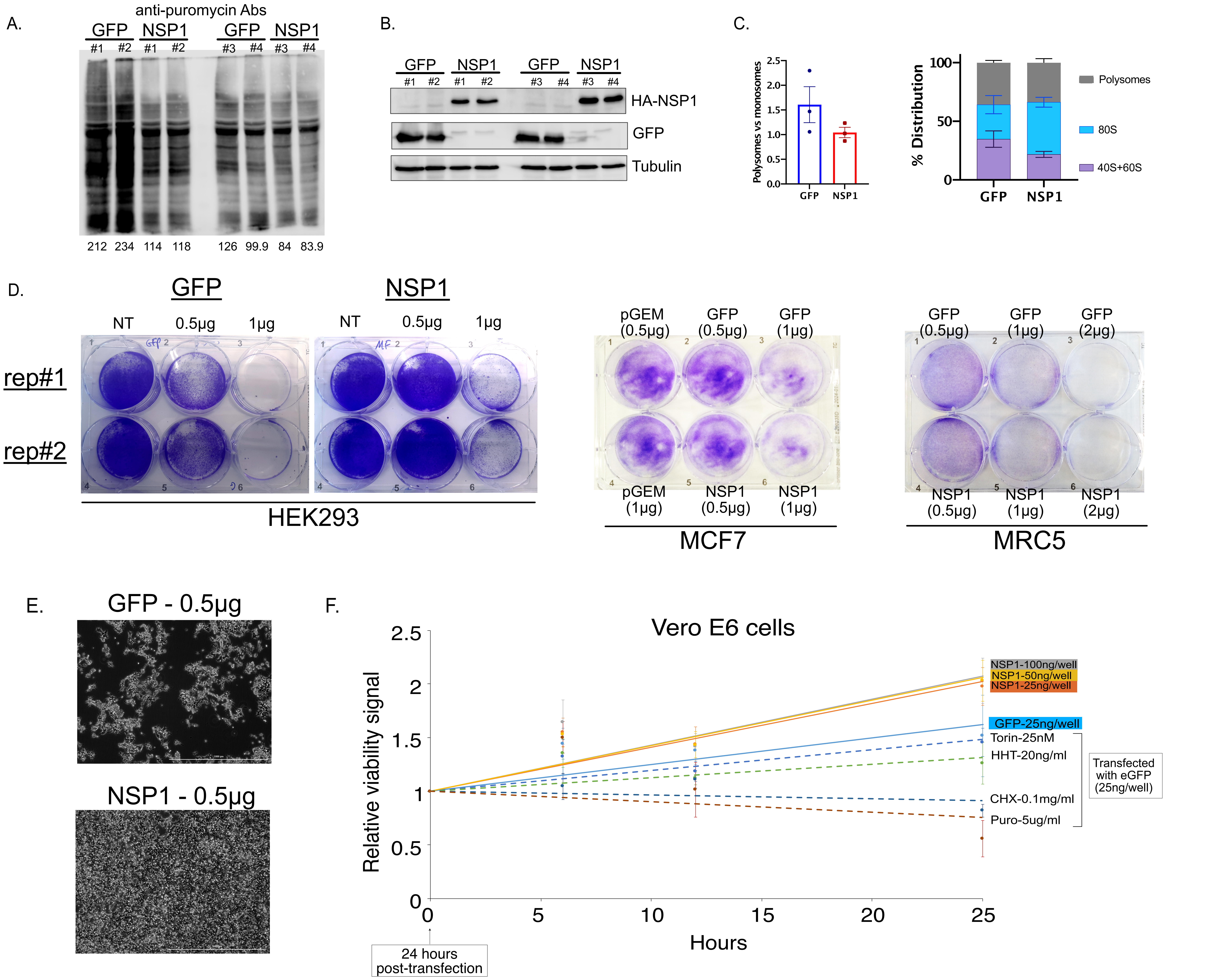


Figure S2: Impact of NSP1 on translation and cell survival. A. Original images of the puromycin labeling experiments presented in Fig. 2A showing probing with anti-puromycin antibodies. B. Western blot analysis of protein expression from the experiments presented on Fig. S1A. In both (A) and (B), four independent experimental repeats are shown. C. Polysomal profiling of MCF7 cells; the experiment was performed as detailed in Fig. 2A. The graphs on the right show the RNA distribution in the free fractions, n=2, bars show STD. D. Indicated cell lines were seeded in 6-wells plates and transfected with plasmids encoding either EGFP or NSP1. After 4 hours of transfection, the medium was changed and the cells were grown for either 72 (for HEK293 cells) or 48 (for other cell lines) hours post-transfection, fixed and stained using violet stain. E. Phase contrast microscopy images of HEK293 cells shown in (D). F. Vero cells were treated as detailed in Fig. 2E and their growth kinetics were detected at the indicated time points; n=3.

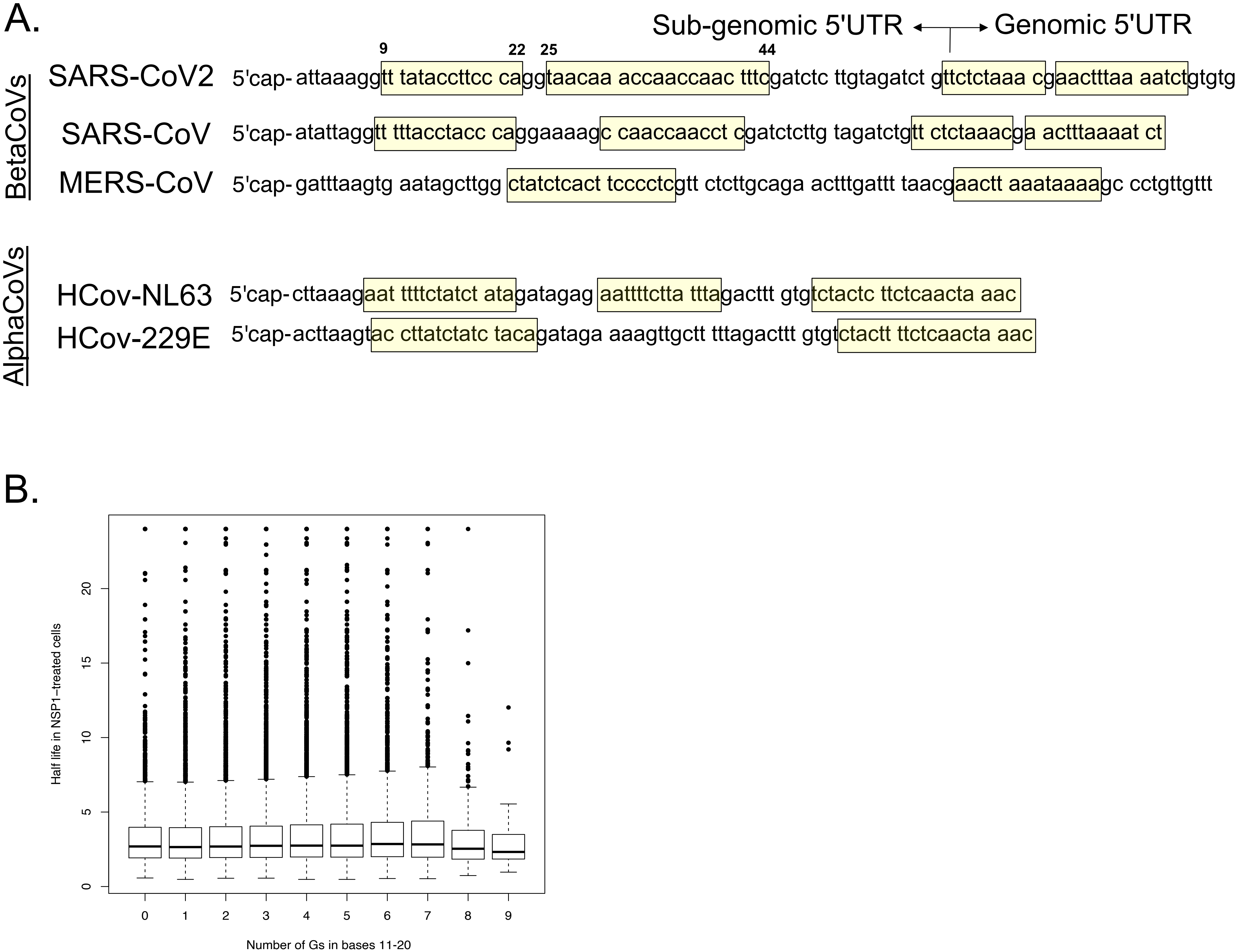


Figure S3: Self-recognition RNA element. A. Guanosine-deficient stretches (yellow boxes) in the cap-proximal regions of various coronavirus-derived 5’UTRs. B. Experiment presented in Fig. 1B was re-analysed to reflect the impact of guanosine residues between nucleotides 11-20 relative to the cap structure on the mRNA stability in the presence of NSP1; n=2.

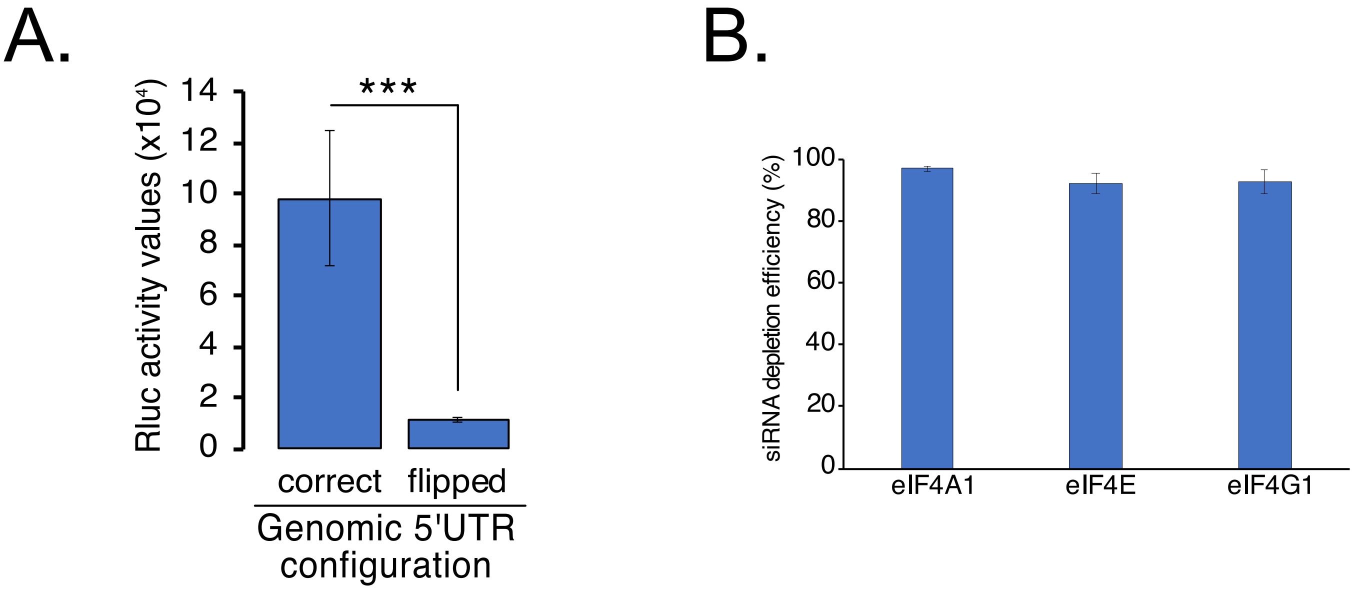


Figure S4. A. MRC5 cells were transfected with in-vitro-transcribed uncapped Rluc mRNAs preceded by SARS-CoV2-derived 5'UTRs introduced in the correct or flipped configuration. Rluc activity was assayed 7 hours after the beginning of transfection; n=4, bars represent SE. B. Knock-down efficiencies of the applied siRNAs. RNA extracted from transfected cells was subjected to qRT-PCR analysis using primers targeting the respective genes and normalized to the expression levels of GAPDH. Reduction in mRNA abundance represents siRNA efficiency;  n=3, bars represent SD.
